# Metaproteomics boosted up by untargeted data-independent acquisition data analysis framework

**DOI:** 10.1101/2020.12.21.423800

**Authors:** Sami Pietilä, Tomi Suomi, Laura L. Elo

## Abstract

Mass spectrometry based metaproteomics is a relatively new field of research that provides the ability to characterize the functionality of microbiota. Recently, we were the first to demonstrate the applicability of data-independent acquisition (DIA) mass spectrometry to the analysis of complex metaproteomic samples. This allowed us to circumvent many of the drawbacks of the conventionally used data-dependent acquisition (DDA) mass spectrometry, mainly the limited reproducibility when analyzing samples with complex microbial composition. However, the previous method still required additional DDA data on the samples to assist the DIA analysis. Here, we introduce, for the first time, a DIA metaproteomics approach that does not require any DDA data, but instead replaces a spectral library generated from DDA data with a pseudospectral library generated directly from the metaproteomics DIA samples. We demonstrate that using the new DIA-only approach, we can achieve higher peptide yields than with the DDA-assisted approach, while the amount of required mass spectrometry data is reduced to a single DIA run per sample. The new DIA-only metaproteomics approach is implemented as open-source software package *DIAtools 2.0*, which is freely available from DockerHub.

## Introduction

Metaproteomics is a relatively new field of research that aims to characterize all proteins expressed by a community of microorganisms in a complex biological sample ^1^. Its major promise lies in its ability to directly measure the functionality of microbiota, while the more widely used metagenomics captures only the taxonomic composition and functional potential. Therefore, metaproteomics has emerged as an intriguing option, for example, in the study of human gut microbiota functionality in various healthy and disease states ^2, 3^.

To date, the common approach in metaproteomics has been data-dependent acquisition (DDA) mass spectrometry, which is, however, known to have limitations ^4^. For example, only the most intense peptide ions are selected for fragmentation, which leaves the rest of the peptides undetected. This is particularly important for metaproteomics, where the vast number of peptides increase the chance of co-elution and peptides are discarded by the instrument from subsequent analysis. The selection also introduces stochasticity to the identifications, reducing the overlap between repeated analyses. For this reason, DDA often requires multiple runs from the same sample to discover all obtainable peptides. Furthermore, the ion intensities are not consistently recorded through the whole chromatographic profile, making quantification challenging ^5^.

To overcome the limitations of DDA, data-independent acquisition (DIA) mass spectrometry produces records of all precursor and fragment ion spectra by systematically fragmenting all precursor peptide ions. Therefore, DIA has been proposed as an alternative method to overcome many fallbacks of DDA. However, the systematic fragmentation of the precursor peptide ions produces highly convoluted fragment spectra, making the peptide identification a difficult task. This is especially challenging for complex metaproteomic samples, where multiple precursor ions are more likely to elute simultaneously.

Recently, we were the first to demonstrate that DIA mass spectrometry can be successfully applied to analyse complex metaproteomic samples by using a spectral library constructed from corresponding DDA data to assist the peptide identification ^6^.

While such DDA-assisted method requires that the peptides are previously discovered through DDA, it allows reproducible identification and quantification of the detected peptides across the samples ^7^. However, the requirement for having a DDA-based spectral library can be considered as a major drawback of the method. Creating the DDA-based spectral library consumes sample material, may not represent well the content of all samples and, most importantly, brings the DDA originated limitations of peptide identification to DIA, as only peptides present in the library can be detected from the DIA data.

Here we introduce, for the first time, untargeted analysis of DIA metaproteomics data without the need for corresponding DDA data. This is done by generating a pseudospectral library directly from the DIA data, following a similar strategy as previously introduced in single-species studies ^8^. The pseudospectra are generated by deconvoluting DIA spectra into DDA-like spectra, having precursors and their fragments, which can then be used for peptide identification with conventional protein database searches. Using laboratory-assembled microbial mixture and human fecal samples, we demonstrate that our DIA-only metaproteomic approach enables overcoming the limitations of the DDA-assisted approach and reduces the number of required mass spectrometry analyses to a single DIA analysis per sample. The new DIA-only metaproteomics approach is implemented as part of our open-source software package *DIAtools 2.0*, which is freely available from DockerHub.

## Results

To demonstrate the feasibility and benefits of our new *DIAtools 2.0* DIA-only metaproteomics approach, we applied it to a laboratory assembled microbial mixture containing twelve different bacterial strains (12mix) and to human fecal samples from six healthy donors (**Supplementary Tables 1-2**) and compared the performance against the previously introduced DDA-assisted method ^6^.

### Peptide identifications and their reproducibility

When investigating the peptide yields, the DIA-only approach of *DIAtools 2.0* produced 13% and 37% higher yields than the DDA-assisted approach in the 12mix (17255 vs. 15242 peptides) and human fecal data (15262 vs. 11136 peptides), respectively (**Figure 1A-B**). Of all the peptides identified by either of the approaches, 88% and 29% were shared between the approaches in the 12mix and the fecal data, respectively, while several peptides were only detected with one of the approaches.

**Figure 1.**
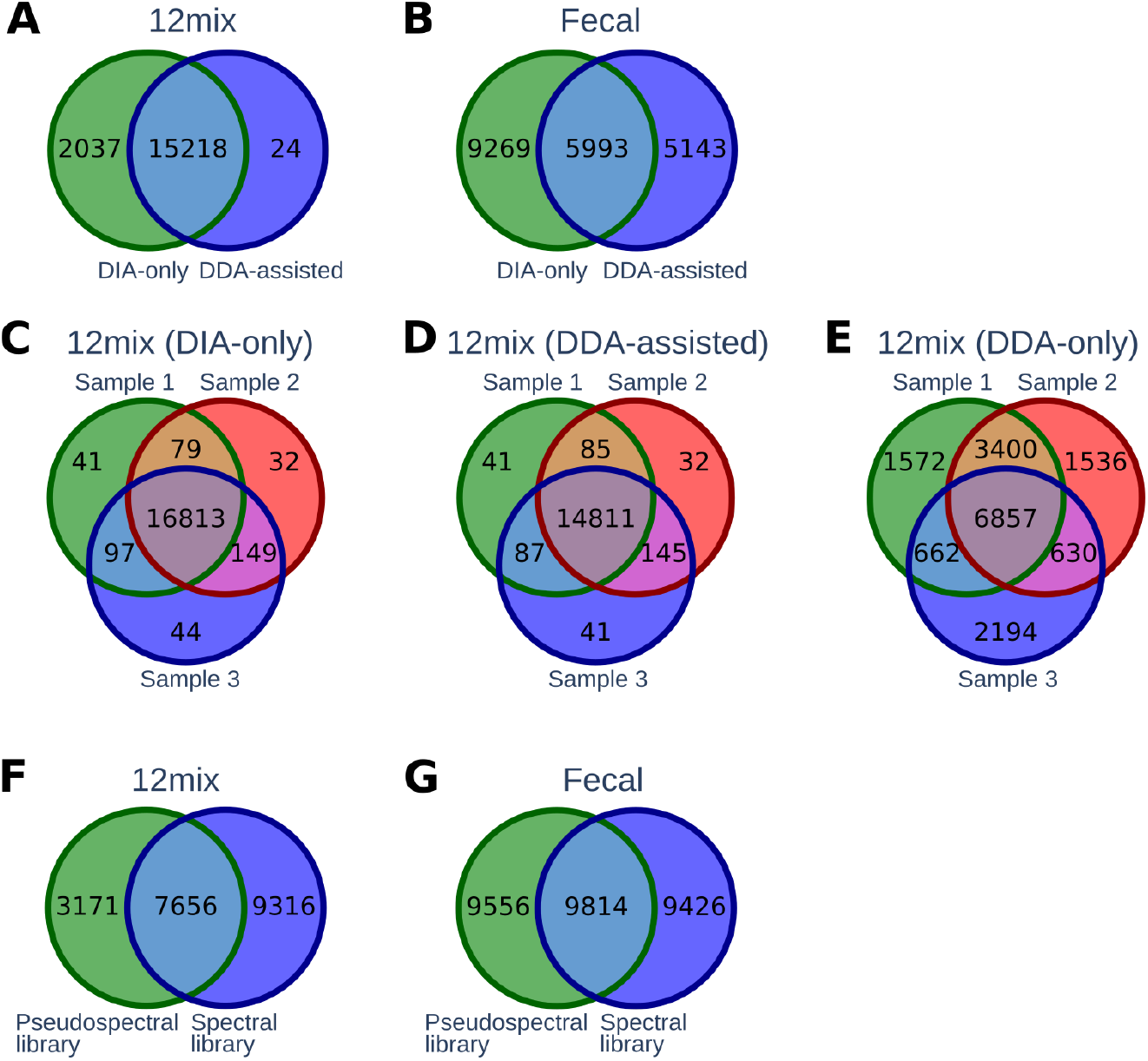
(**A-B**) Overlap of peptides detected by the *DIAtools 2.0* DIA-only and the DDA-assisted approach in the 12mix and the human fecal datasets. (**C-E**) Overlap of detected peptides between three replicated 12mix samples using the DIA-only, DDA-assisted, and DDA-only approach. (**F-G**) Overlap of peptides between the spectral and pseudospectral libraries in the 12mix and the human fecal datasets.

In the 12mix data, we also investigated the reproducibility of the identified peptides between three technical replicates. The reproducibility of both the DIA-only and the DDA-assisted approach was high, with an overlap of over 97% in both cases (**Figure 1C-D**). For comparison, use of only the DDA data resulted in an overlap of only 41% (**Figure 1E**), highlighting the improved reproducibility of DIA over DDA.

Next, we explored the composition of the spectral and pseudospectral libraries, which is an essential technical aspect of the peptide identification, forming the search space of peptides that can be detected from the DIA data. Of all the library peptides, 38% and 34% were shared between the spectral and pseudospectral libraries in the 12mix and the human fecal data, respectively (**Figure 1F-G**). Interestingly, although in the 12mix data, the DDA-based spectral library contained more peptides than the DIA-based pseudospectral library, the latter resulted in more peptide identifications (**Figure 1A**). This implies that, as expected, a library built directly from the DIA data targets the DIA data peptides more precisely than a separate DDA-based library.

### Taxonomic and functional profiles

Overall, the taxonomic and functional profiles of the metaproteomes observed using the *DIAtools 2.0* DIA-only and the DDA-assisted approaches were highly similar. The DIA-only approach was able to assign a unique taxonomic annotation at genus level to 60% of the peptides in the 12mix (**Figure 2A**) and 43% of the peptides in the human fecal data (**Figure 2B**). Similarly, the DDA-assisted approach annotated 61% and 41% of the peptides in the 12mix and human fecal data, respectively (**Figure 2A-B**). With both the DIA-only and the DDA-assisted approach, less than 2% of the identified peptides in the laboratory assembled 12mix were annotated to genera not present in the mixture. The taxonomic profile of human fecal samples was similar to those reported in the literature ^2, 9^. These provide further validation of the feasibility of DIA metaproteomics in profiling complex microbial samples.

**Figure 2.**
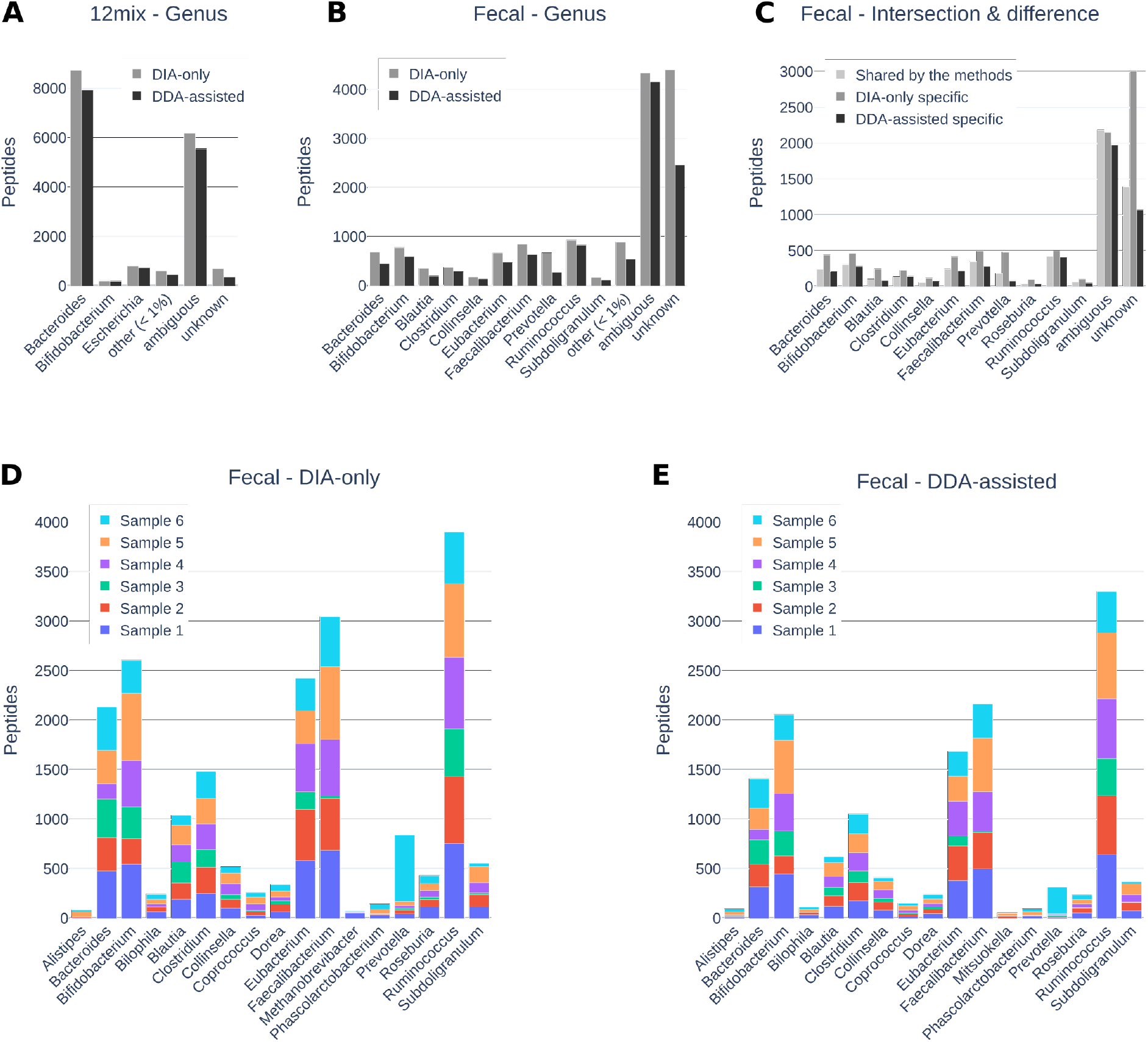
Genus-level taxonomic profiles of the (**A**) 12mix and (**B-C**) human fecal samples using the *DIAtools 2.0* DIA-only or the DDA-assisted approach. Genera having less than 1% of the total peptides were aggregated to category *other*. Genus-level taxonomic profiles of the individual human fecal samples using (**D**) the DIA-only or (**E**) the DDA-assisted approach.

To understand better the differences between the DIA-only and DDA-assisted peptide identifications, we compared the genus-level taxonomic profiles of the peptides found only by one of the approaches (**Figure 2C**). While the overall taxonomic profiles of these peptides were highly similar, some notable differences were observed. First, while the DIA-only approach detected a larger number of peptides, it also detected a larger proportion of unknown peptides with no taxonomic annotation in the widely used integrated reference catalog of the human gut microbiome (IGC) ^10^. This suggests that the most abundant proteins, commonly targeted by DDA, are more likely to have an annotation in the database. Secondly, the DDA-assisted method detected a larger proportion of ambiguous peptides with multiple different annotations in IGC, which is typical to peptides that are shared by multiple organisms. Again, this is in line with DDA targeting the most abundant peptides. Finally, when investigating the individual genera, the largest difference between the results of the DIA-only and DDA-assisted approaches was observed in *Prevotella* in the human fecal samples, which had a 4% share with the DIA-only approach and 2% share with the DDA-assisted approach. A closer look at the individual human fecal samples revealed that *Prevotella* was dominant in a single sample, while its proportion in the other samples was very low (**Figure 2D-E**). This highlights the limitations of using a pooled DDA library for the analysis, which may adversely affect the performance of the DDA-assisted approach.

Similarly, we investigated the identified KEGG functional profiles of the peptides and found them to be highly similar between the DIA-only and DDA-assisted approaches (**Supplementary Figure 1**).

### Peptide quantifications and their reproducibility

Finally, we assessed the performance of the *DIAtools 2.0* DIA-only and the DDA-assisted approach in quantifying the identified peptides. With both approaches, the pairwise Pearson correlation coefficients between the quantifications across the technical replicates in the 12mix data were very high (r > 0.95 with p < 0.001 in each pairwise comparison, **Figure 3A-B** and **Supplementary Figure 2**), indicating high reproducibility of the approaches.

**Figure 3.**
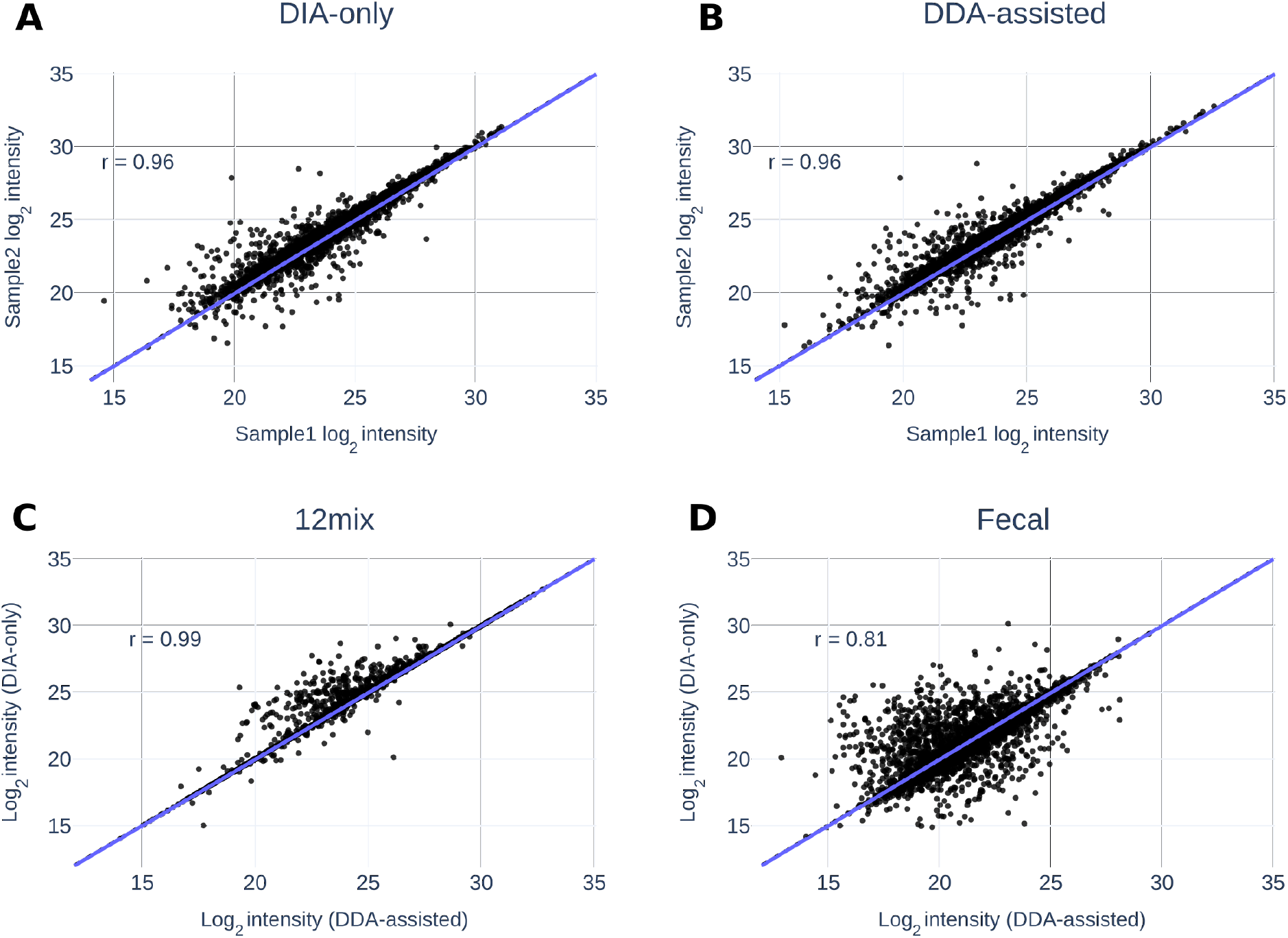
Representative examples of correlations of peptide quantifications between two technical replicates of the 12mix samples using (**A**) the DIA-only or (**B**) the DDA-assisted approach. All the corresponding pairwise comparisons are shown in **Supplementary Figure 2.** Correlations of peptide quantifications between the DIA-only and the DDA-assisted approach in (**C**) the 12mix and (**D**) the human fecal samples.

Further investigation of the quantifications by the DIA-only and the DDA-assisted approaches suggested both similarities and differences. The overall correlations between the approaches were high: 0.99 for the 12mix (p < 0.001) and 0.81 for the human fecal samples (p < 0.001) (**Figure 3C-D**). In general, however, the DIA-only estimates for many of the peptides were somewhat higher than the estimates produced by the DDA-assisted approach. This is related to differences in the libraries built by the approaches, where each unique peptide can be represented by multiple ions with different charge states and modifications and the same peptide is typically represented by a partially different set of fragments (**Supplementary Figure 3**). This results in quantification differences between the approaches, because the peptides are quantified on the basis of the fragment level.

## Discussion

We introduce here a new computational tool for DIA metaproteomics to overcome the main limitations of the currently used DDA-based methods. Our results suggest that our new DIA-only approach of *DIAtools 2.0* enables reproducible identification and quantification of metaproteome profiles beyond both the currently used DDA-assisted DIA approach or the more widely used DDA-only method. In particular, the *DIAtools 2.0* DIA-only approach improves both the number of detections as well as their reproducibility over the DDA-based methods. Another major benefit of the proposed approach is that it reduces the number of samples that need to be analysed, as additional DDA-based analysis is no longer needed for library generation. This is of particular importance in studies with large numbers of samples, such as those in clinical study settings.

Microbiome profiling has attained increasing attention in the past few years with the recognition of the important role of microbiota in human health and disease. For instance, gut microbiome has been associated with various health and disease states and has been suggested as the *“forgotten”* human organ ^11–13^. Overall, better understanding of how the complex microbial communities in human influence disease pathogenesis has major implications for disease prediction, prevention, and treatment.

Currently, the most common approach to study microbiome is metagenomics. It has been successfully applied in various studies, including large multi-centre studies of thousands of samples using either 16S rRNA or whole genome sequencing ^14^. By cataloguing which microbes are present in a sample and their relative abundances, metagenomics can provide important information about the taxonomic composition of the microbial communities and predict their functional potential. A major limitation of the metagenome approach is, however, that it does not directly assess the function of the microbiota. To overcome this limitation, mass spectrometry based metaproteome analysis has emerged as an alternative option.

In addition to technical challenges, a major bottleneck in the utilisation of mass spectrometry metaproteomics is the lack of appropriate computational tools to interpret the data produced ^15^. Because of the inherent complexity of the data, conventional tools to analyse single-species proteome data are often not well suited for metaproteomics. While a few tools have been introduced for the analysis of DDA metaproteome samples ^16,17^, until our recent work ^6^, there have been no tools for DIA mass spectrometry metaproteomics, despite its high potential to improve the reproducibility over the more conventionally used DDA mode. The aim of this study was to address that need.

To this end, we propose here a major practical improvement to DIA metaproteomics analysis by showing, for the first time, that reproducible identification and quantification of microbial peptides is possible without the need for additional DDA library samples using our new DIA-only *DIAtools 2.0*. In addition to a reduced number of samples that need to be analysed, this circumvents the initial DDA-originated limitations in peptide identification that hamper approaches using a DDA-based spectral library, especially as such library is typically prepared using only a limited set of pooled samples. This may play a crucial role in the data interpretation, as demonstrated in the analysis of the human fecal samples, where *Prevotella* was dominant in one of the samples but lowly abundant in the others. If the DDA library does not well represent the whole sample set, important parts of the microbial communities may remain undetected, which can be avoided with the DIA-only approach.

The peptide yields from the DIA metaproteomics methods were comparable to those reported in DDA metaproteomic studies with similar laboratory protocols ^18^. Importantly, for the DIA-only approach this was achieved with only a single analysis run per sample. Notably, the majority of the peptides detected using the DDA-assisted DIA approach were detected also using the DIA-only method, whereas the total number of peptides detected by the DIA-only method exceeded the number of peptides detected by the DDA-assisted method. This confirms that a library built from DDA data is not required for DIA metaproteomics.

A well-known limitation of the DDA-based analysis is that it tends to detect peptides that are highly abundant. This includes peptides shared by multiple species, which was observed here as the relatively large proportion of peptides with ambiguous taxonomic annotation with the DDA-assisted approach. On the other hand, the DIA-only approach tended to detect a larger proportion of peptides with unknown genus, indicating that they were less well characterized in the current databases. This may suggest ability to target proteins not detectable by other means, thus revealing proteins that are not well known and annotated by databases.

Technically, analysis of metaproteomics data can be a very computationally intensive task in terms of the required computing power and memory usage. For building a spectral or pseudospectral library, a great deal of computing power is needed to compare the theoretical spectra of the vast sequence database against the experimental spectra obtained from the DIA data. With the datasets in this study, this was found to be the most time-consuming step that can take several days. Once the spectral or pseudospectral library has been produced, the subsequent analysis of the DIA data against the library spectra is then considerably faster. The current version of *DIAtools 2.0* scales the processing with threads using efficiently the processing power of a single computer. However, it is possible to extend the parallel analysis to multiple computers, such as cluster environments. To enable easy deployment of *DIAtools 2.0*, it is implemented as a software container which provides all the required utilities and libraries in a single package.

An interesting future development would be to circumvent the need of generating reference spectra separately for each new project. For this, machine learning has been suggested as a possible solution using, for instance, artificial neural networks ^19–21^. A major challenge with such approaches is, however, their potential biases towards the training data and need for re-training for specific conditions. This remains an interesting topic for further investigation.

Overall, the microbiome research still involves multiple different types of unknowns that are continuously being revealed thanks to improved technologies ^22^. In this endeavour, metaproteomics provides an excellent opportunity to uncover the functionally important aspects of the microbial communities, providing complementary information to the studies of microbiomes and their role in health and disease.

## Materials and methods

### Generation of DIA-based pseudospectral library

To enable building a pseudospectral library directly from the DIA data, the spectra in the DIA data were deconvoluted into pseudospectra containing precursor ions and their corresponding fragment spectra following a similar procedure as in the DIA-Umpire tool for single-species proteomics ^8^. In short, a 2D feature detection algorithm was first used to locate precursor and fragment ions from the MS1 and MS2 data. Pearson correlation coefficients of the elution peaks and retention time differences of the peak apexes were then used to pair precursor and fragment ions. For the generation of pseudospectra, all likely complementary *y* and *b* ions were detected. The obtained DIA pseudospectra were finally searched with X!Tandem ^23^ and Comet ^24^ algorithms against the Integrated reference catalog of the human gut microbiome (IGC, 9.9M) ^10^, containing over 9 million protein sequences covering human gut bacteria. The false discovery rate (FDR) for the identifications was set at 1%. The identified spectra formed the final pseudospectral library that was used to identify peptides from the DIA data.

### Peptide identification and quantification

Peptide identifications and quantifications were obtained from the DIA data using *DIAtools 2.0*. For peptide identification, either the DIA-based pseudospectral library (referred to as DIA-only approach) or the DDA-based spectral library (referred to as DDA-assisted approach) was used on the basis of X!Tandem ^23^ and Comet ^24^ algorithms and the IGC reference database. Parent ion mass tolerance was set to 10 ppm and fragment ion tolerance to 0.02 Da. The false discovery rate (FDR) for the spectral library matching was set at 1%. For TRIC feature alignment ^25^, the target and maximum FDRs were set to 1% and 5%, respectively.

### Taxonomic and functional annotations

The identified peptides were taxonomically and functionally annotated using the annotations from the IGC database without protein inference. For each peptide, annotations of all possible protein sequences were retrieved. An annotation was assigned to a peptide only if there was no evidence of conflicting annotations. In case of conflicting annotations, a peptide was annotated as ambiguous.

### Software and availability

The *DIAtools 2.0* software is open source and distributed as a Docker image that can be downloaded from DockerHub repository elolab/DIAtools-2.0. The image is based on Ubuntu 18.04 and comes bundled with several programs and libraries that retain their original licenses. The installed software include: Comet 2017.01 rev. 4, X!Tandem 2017.02.01.4, OpenMS 2.4 (includes OpenSWATH), Trans-Proteomic Pipeline (TPP) 5.1, msproteomicstools 0.8.0, SWATH2stats 1.8.1, DIA-Umpire 2.1.3, and Thermo msFileReader. The source code and step-by-step instructions to use the software are provided at https://github.com/elolab/DIAtools.

### Laboratory assembled microbial mixture and human fecal samples

The 12mix data was a mixture of twelve (12) different bacterial strains isolated from fecal samples of three human donors grown on fastidious anaerobe agar (LAB 090; LAB M, UK) and annotated by sequencing their 16S-rDNA: *Bacteroides vulgatus*, *Parabacteroides distasonis*, *Enterorhabdus sp.*, *Bifidobacterium pseudocatenulatum*, *Escherichia coli*, *Streptococcus agalactiae*, *Bacteroides fragilis*, *Alistipes onderdonkii*, *Collinsella aerofaciens*, *Clostridium sordellii*, *Eubacterium tenue*, and *Bifidobacterium bifidum*. Prior to mixing, the bacterial cell counts were equalized to 10 x 10^8^ cells / ml using flow cytometry and 1 x 10^8^ cells of each isolate were added to the final mixture. Three isolations and mixtures were made, and each mixture was analyzed in DDA and DIA mode on a Q Exactive HF mass spectrometer (Thermo Fisher Scientific) equipped with a nano-electrospray ionization source, as described in Aakko *et al.* (2019) ^6^.

The human fecal data contained six human fecal samples from anonymous individuals representing a complex metaproteomic scenario. Each fecal sample was analyzed in DIA mode with a single injection. Additionally, all six samples were pooled together and analyzed in DDA mode with six injections. The analyses were performed on a nanoflow HPLC system (Easy-nLC1200, Thermo Fisher Scientific, Waltham, Massachusetts, USA) coupled to a Q Exactive HF mass spectrometer (Thermo Fisher Scientific) equipped with a nano-electrospray ionization source, as described in Aakko *et al.* (2019) ^6^.

The mass spectrometry data are available from the ProteomeXchange Consortium via the PRIDE partner repository with the dataset identifier PXD008738.

## Acknowledgements

The authors wish to acknowledge Dr. Juhani Aakko for his insights during the conception of the study. Prof. Elo reports grants from the European Research Council ERC (677943), Academy of Finland (296801, 304995, 310561, 314443, and 329278), and Sigrid Juselius Foundation, during the conduct of the study. Our research is also supported by University of Turku Graduate School (UTUGS), Biocenter Finland, and ELIXIR Finland.

## Author contributions

LLE conceived and supervised the study. SP designed the data analysis approach, implemented the software package and performed the data analysis. SP, TS and LLE interpreted the results, and wrote the manuscript. All authors have read the final version of the manuscript and approved of its content.

## Code Availability Statement

All source code generated for this work will be released under open source license (GNU General Public License v3.0) and will be stored in a public repository (https://github.com/elolab/DIAtools).

