## Supplementary Material for "Metaproteomics boosted up by untargeted data-independent acquisition data analysis framework"

Sami Pietilä[1,*], Tomi Suomi[1,*], Laura L. Elo[1,2,#]

[1] Turku Bioscience Centre, University of Turku and Åbo Akademi University, FI-20520 Turku, Finland

[2] Institute of Biomedicine, University of Turku, FI-20520 Turku, Finland

* Shared first author

[#]

### Table of contents

**S-1** Supplementary Figure 1. KEGG functional profiles of the 12mix and human fecal samples using the DIA-only or the DDA-assisted metaproteomics approach.

**S-2** Supplementary Figure 2. Correlations of peptide quantifications between each pairwise comparison of technical replicates of the 12mix samples using the DIA-only or the DDA-assisted approach.

**S-3** Supplementary Figure 3. The proportion of peptide ion spectra in the library with different numbers of matching fragment ions between the DIA-only and the DDA-assisted approach in the 12mix and human fecal datasets.

**S-3** Supplementary Table 1. 12mix dataset files.

**S-4** Supplementary Table 2. Human fecal dataset files.

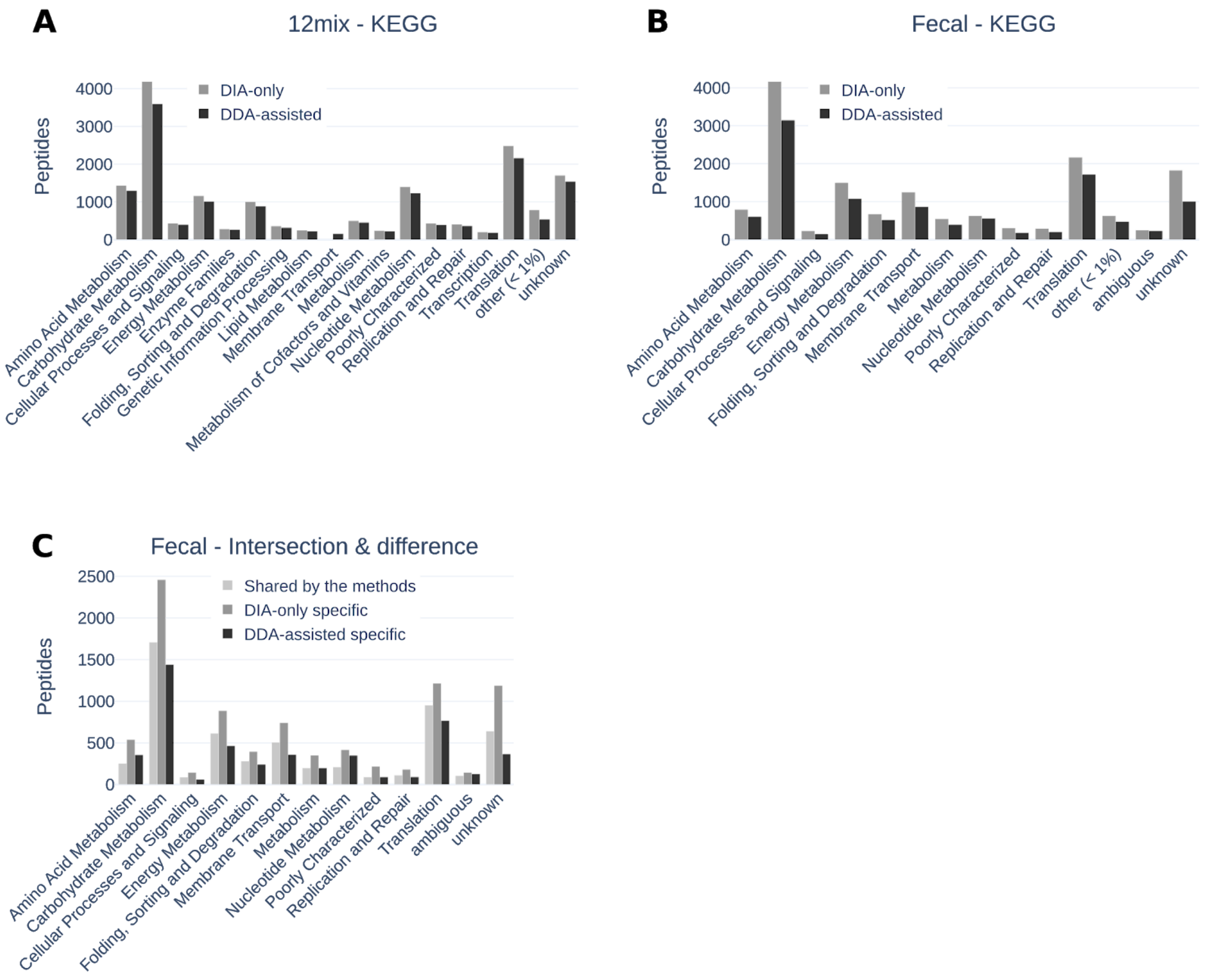


**Supplemental Figure 1.** KEGG functional profiles of the **(A)** 12mix and **(B)** human fecal samples using the DIA-only or the DDA-assisted metaproteomics approach. KEGG functions with less than 1% of total peptides were aggregated in the figure to the category 'other'. **(C)** KEGG functional profiles of the peptides detected by both the DIA-only and the DDA-assisted approaches, or by only one of the approaches in the human fecal data.

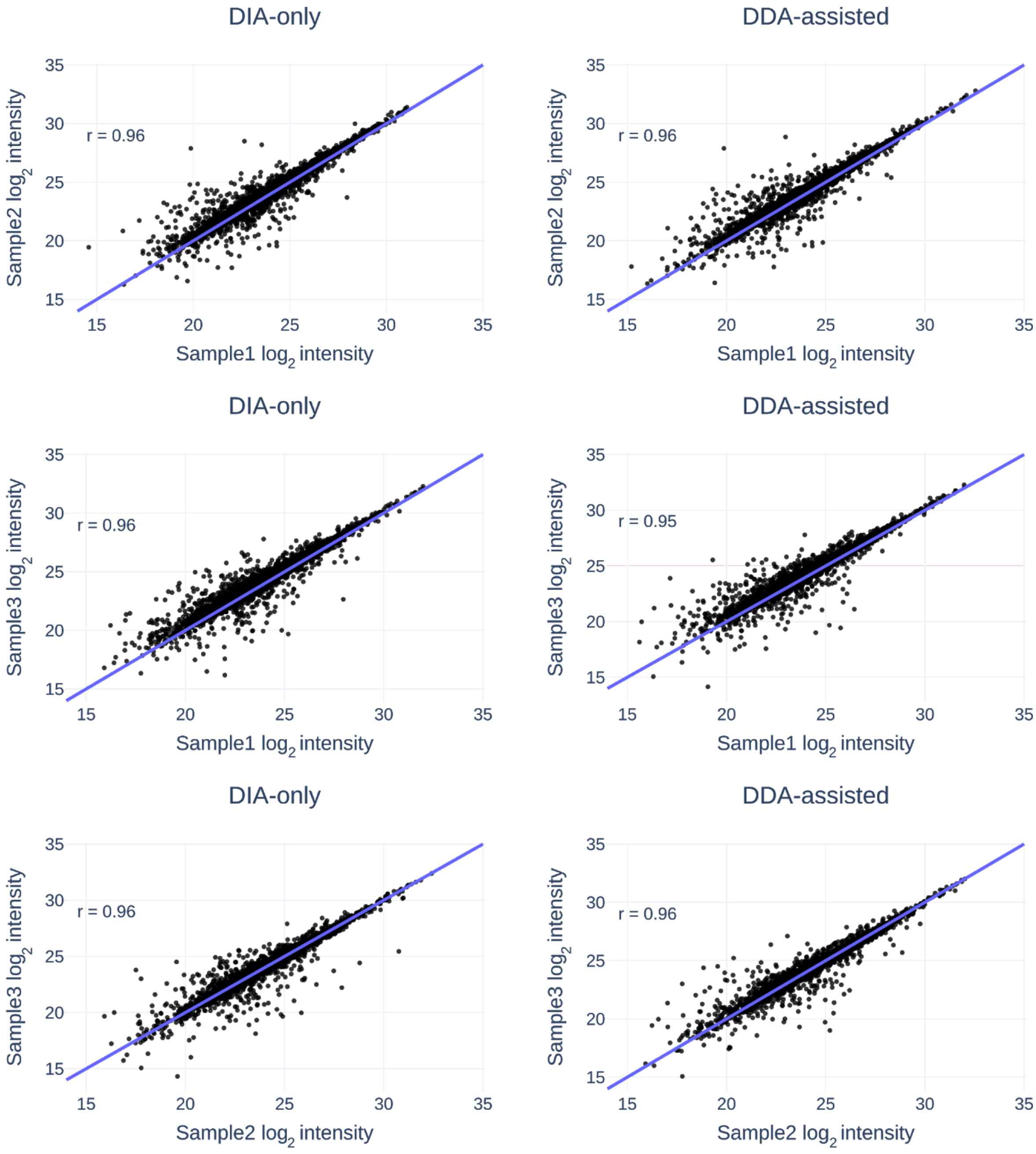


**Supplemental Figure 2.** Correlations of peptide quantifications between each pairwise comparison of technical replicates of the 12mix samples (rows) using the DIA-only or the DDA-assisted approach (columns). All Pearson correlation coefficients (r) were highly significant $p < 0.001$.

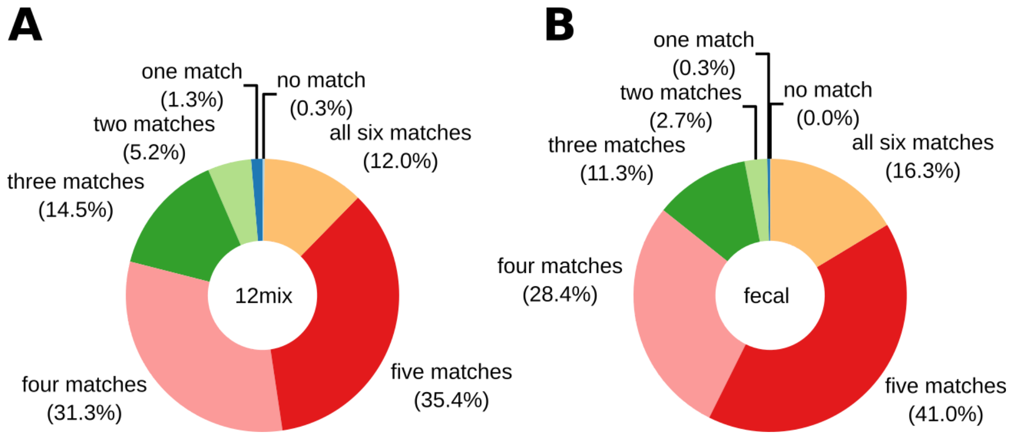


**Supplementary Figure 3.** The proportion of peptide ion spectra in the library with different numbers of matching fragment ions between the DIA-only and the DDA-assisted approach in the **(A)** 12mix and **(B)** human fecal datasets. The library was built in such a way that each peptide spectrum was represented with one precursor ion and exactly six fragment ions.

**Supplementary Table 1.** 12mix dataset files. The files have been deposited to the ProteomeXchange Consortium via the PRIDE partner repository with the dataset identifier, PXD008738.

| **File Type** | **File Name** | **Description** |
|---|---|---|
| DDA | 170412_12mix_DDA_library_stock_2_2.raw | Pooled spectra for library |
| DDA | 170412_12mix_DDA_library_stock_2_3.raw | Pooled spectra for library |
| DDA | 170412_12mix_DDA_library_stock_2_4.raw | Pooled spectra for library |
| DIA | 170413_12mix_DIA_14.raw | Replicated Sample #1 |
| DIA | 170413_12mix_DIA_15.raw | Replicated Sample #2 |
| DIA | 170413_12mix_DIA_16.raw | Replicated Sample #3 |

**Supplementary Table 2.** The human fecal dataset files. The files have been deposited to the ProteomeXchange Consortium via the PRIDE partner repository with the dataset identifier PXD008738.

| File Type | File Name | Description |
|---|---|---|
| DDA | 170825_HF_1_6_pool_2ug_DDA_library_1column_1.raw | Pooled spectra for library |
| DDA | 170825_HF_1_6_pool_2ug_DDA_library_1column_2.raw | Pooled spectra for library |
| DDA | 170825_HF_1_6_pool_2ug_DDA_library_1column_3.raw | Pooled spectra for library |
| DDA | 170825_HF_1_6_pool_2ug_DDA_library_1column_4.raw | Pooled spectra for library |
| DDA | 170825_HF_1_6_pool_2ug_DDA_library_1column_5.raw | Pooled spectra for library |
| DDA | 170825_HF_1_6_pool_2ug_DDA_library_1column_6.raw | Pooled spectra for library |
| DIA | 170825_HF_1ug_DIA_1column_1.raw | Sample 1 |
| DIA | 170825_HF_1ug_DIA_1column_2.raw | Sample 2 |
| DIA | 170825_HF_1ug_DIA_1column_3.raw | Sample 3 |
| DIA | 170825_HF_1ug_DIA_1column_4.raw | Sample 4 |
| DIA | 170825_HF_1ug_DIA_1column_5.raw | Sample 5 |
| DIA | 170825_HF_1ug_DIA_1column_6.raw | Sample 6 |
